## Supplemental Figures and Tables for "Cryo-ET distinguishes platelets during pre-acute myeloid leukemia from steady state hematopoiesis"

**Figure S1. Segmentation, training and measurement of platelet and organelles.**

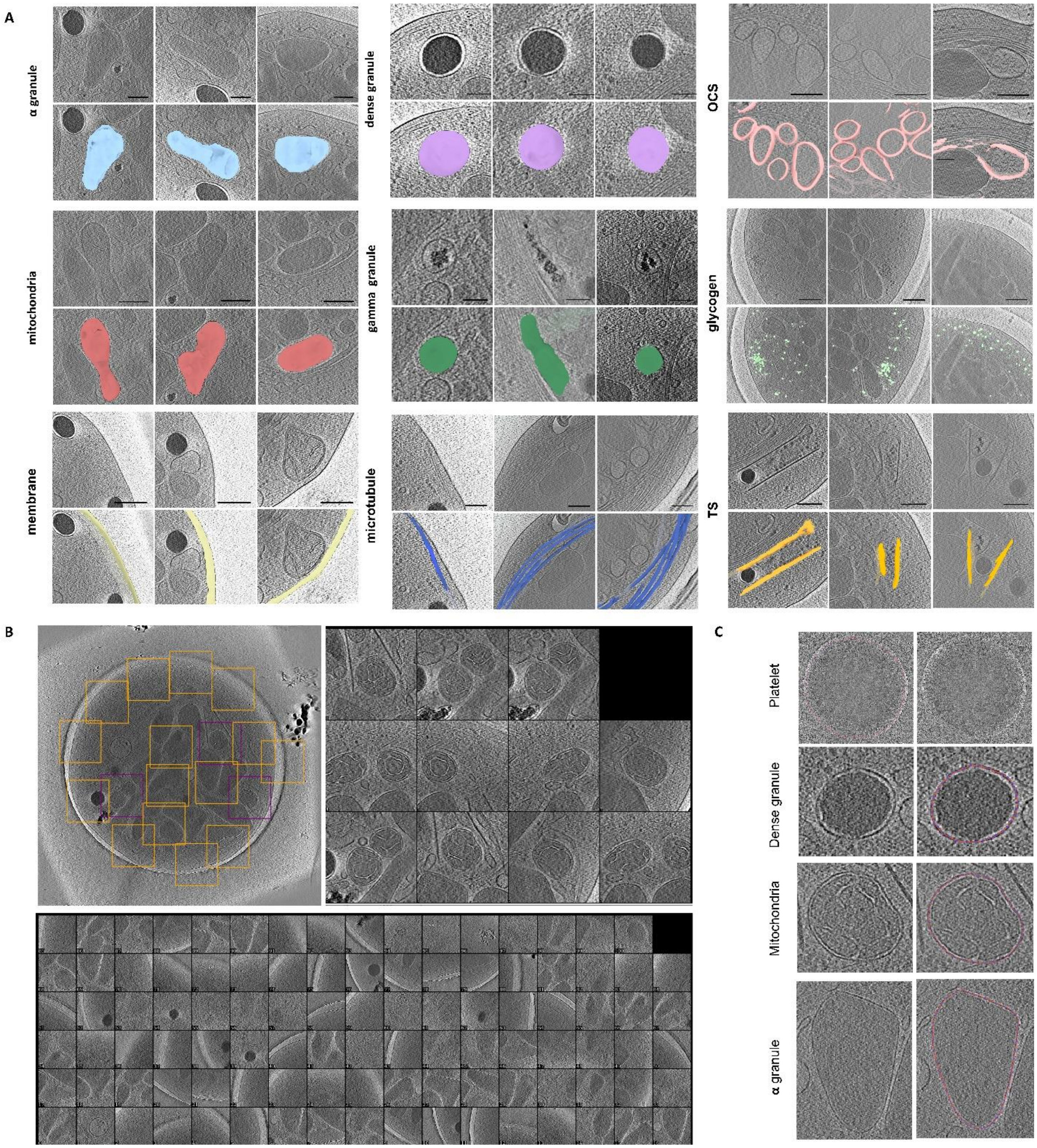


A, Segmentation of organelles in platelet: α granule (scale bar 200 nm), dense granule (scale bar 100 nm), mitochondria (scale bar 300 nm), gamma granule (scale bar 100 nm), membrane (scale bar 300 nm), microtubule (scale bar 200 nm), open canalicular system (OCS) (scale bar 200 nm), glycogen (scale bar 300 nm), tubule-like system (TS) (scale bar 200 nm). B, Training set of mitochondria. A slice of tomogram to pick the positive (purple box) and negative control (yellow box) for mitochondria. C, Area measurement.

**Figure S2 Cryo-ET of platelets from Un-irradiated, AML and pre-AML mice.**

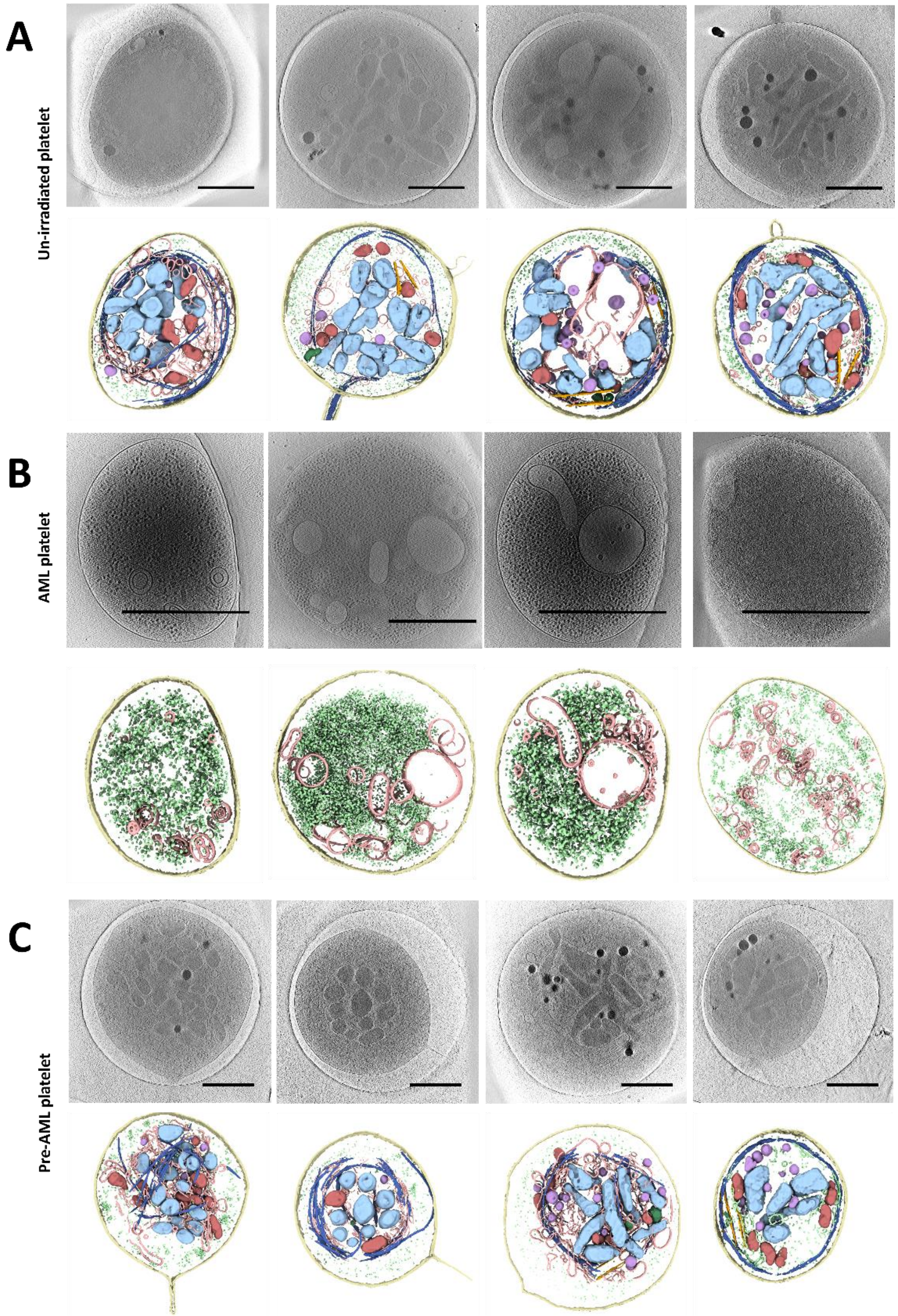


A, Normal platelets had typical features (scale bar 1 µm). B, The platelets of leukemia showed small round or oval shape with only glycogen particles and open canalicular system (OCS) inside. (scale bar 1 µm) C, Pre-AML platelets had α granules, dense granules, λ granules, normal mitochondria, plasma membrane, microtubule, glycogen particles, and OCS (scale bar 1 µm).

**Figure S3. Statistics for alpha granules and dense granules of platelets**.

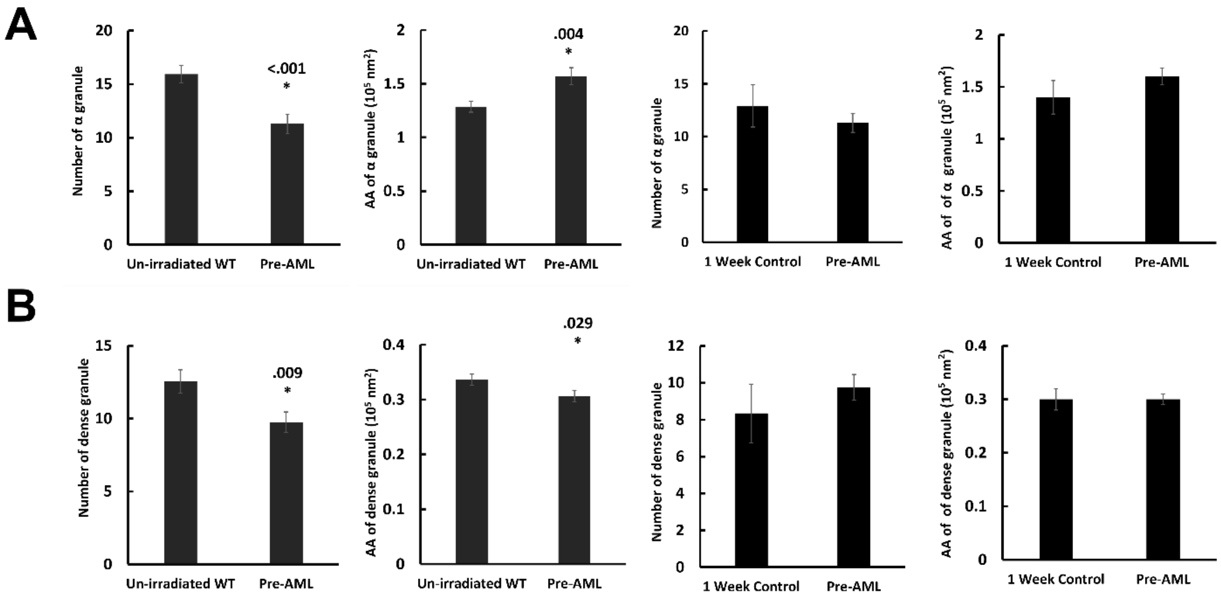


A, Compared with un-irradiated mice, pre-AML platelets increased in the average area (mean area of all α granules in the same platelet cell) of α granule ($P$=0.004) and decreased in the number of α granule ($P$<0.001). (AA: Average area). The number and average area of α granule between 1 week bone marrow transplantation (BMT) control and pre-AML platelets has no significant changes. B, Dense granules in pre-AML platelets decreased in the number ($P$=0.009) and average area (mean area of all dense granules in the same platelet cell) ($P$=0.029), compared with platelets from un-irradiated mice. No significant changes found between 1 week BMT control and pre-AML platelets.

**Figure S4. Mitochondria microenvironment and identification.**

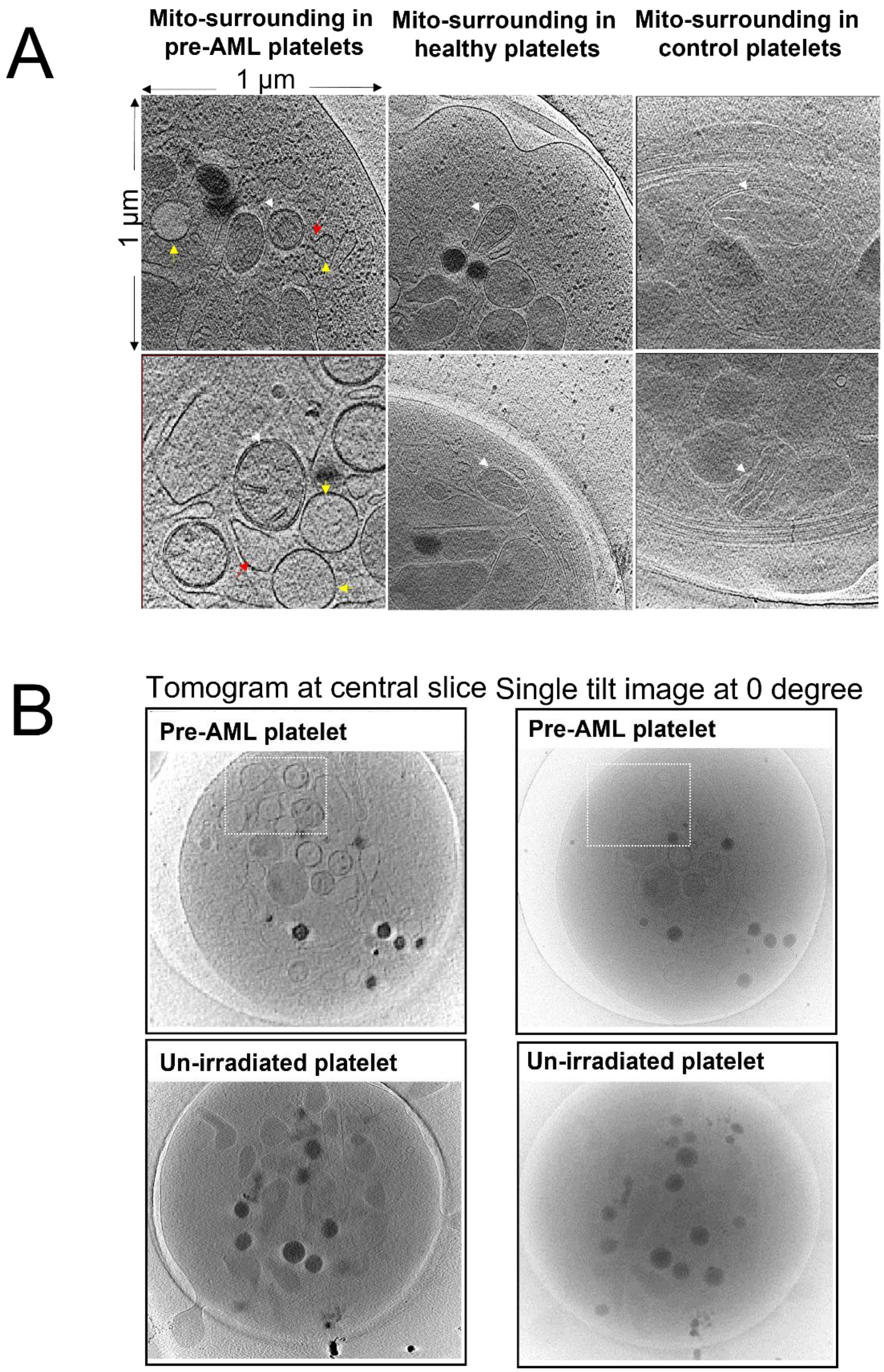


A. Selection of mitochondria microenvironment at single tomogram slices by boxing out 1 $\mu m^2$ area with mitochondria centered. (Upper lane: abnormal mitochondria; Lower lane: normal mitochondria; White arrow: mitochondria; Yellow arrow: spherical vesicles; Red arrow: possible connections) B. Formation of cluster of spherical vesicles (labeled by white rectangle) could be easily seen in reconstructed tomogram and even visible in a single image.

**Supplemental Table 1. Comparison of the area of platelets in different groups**

| | **Un-irradiated (94)** | **Pre-AML (72)** | **AML (110)** | **BMT Control (19)** | **P-value (Un-irradiated vs. Pre-AML)** | **P-value (Un-irradiated vs. AML)** | **P-value (Un-irradiated vs. BMT Control)** |
|---|---|---|---|---|---|---|---|
| Area of platelet ($10^5$ $nm^2$) | 75.6 ± 1.84 | 73.0 ± 2.5 | 15.6 ± 1.0 | 68.4 ± 3.03 | .393 | <.001 | .099 |

Un-irradiated, platelets from wild-type mice; Pre-AML, platelets from mice at the first week after transplantation with leukemia progenitor and stem cells; AML, platelets from mice at the third week after transplantation with leukemia progenitor and stem cells; BMT Control, platelets from mice at the third week after transplantation with normal bone marrow cells.

**Supplemental Table 2. Statistics of α granule, dense granule, and mitochondria**

| | **Un-irradiated (94)** | **Pre-AML (72)** | ***P*-value** |
|---|---|---|---|
| **α granule** | | | |
| Number | 15.9 ± 0.8 | 7.7 ± 0.9 | <.001 |
| Average area ($10^5$ $nm^2$) | 1.3 ± 0.05 | 1.6 ± 0.08 | .004 |
| Area percentage (%) | 24.2 ± 0.7 | 20.7 ± 1.2 | .013 |
| **Dense granule** | | | |

| | | | |
|---|---|---|---|
| Number | 12.5 ± 0.8 | 9.7 ± 0.7 | .009 |
| Average area ($10^5$ $nm^2$) | 0.34 ± 0.01 | 0.31 ± 0.01 | .029 |
| Area percentage (%) | 5.3 ± 0.3 | 4.1 ± 0.3 | .003 |
| **Mitochondria** | | | |
| Number | 8.1 ± 0.5 | 7.0 ± 0.6 | .161 |
| Average area ($10^5$ $nm^2$) | 0.6 ± 0.01 | 0.7 ± 0.03 | .023 |
| Area percentage (%) | 6.5 ± 0.3 | 6.6 ± 0.6 | .826 |

Un-irradiated, platelets from wild-type mice; Pre-AML, platelets from mice at the first week after transplantation with leukemia progenitor and stem cells; Average area, mean area of all objects of interest in the same platelet; Area percentage, total area of all objects of interest to the area of platelet.
